# Activation loop phosphorylation of Cdk11 is restrained by PNUTS-PP1 and regulates Cdk11 activity and function

**DOI:** 10.1101/2024.05.08.592654

**Authors:** Amy E Campbell, Abdulrahman A Aljabri, Andrew Hesketh, Dominic P Byrne, Hayley Bennett, Sanjai Patel, Philip Brownridge, Thomas Zacharchenko, Giselda Bucca, Patrick A Eyers, Andrea J Betancourt, Claire E Eyers, Daimark Bennett

**Affiliations:** Faculty of Health and Life Sciences, University of Liverpool, Biosciences Building, Crown Street, Liverpool, L69 7ZB UK; Kinomica Ltd, Manchester Science Park, Mereside, Macclesfield, SK10 4TG, UK; Faculty of Biology, Medicine and Health, University of Manchester, Michael Smith Building, Manchester, M13 9PT, UK; Department of Pharmacology and Toxicology, College of Pharmacy, Taibah University, Madinah, Kingdom of Saudi Arabia; School of Applied Sciences, University of Brighton, Cockcroft Building, Lewes Road, Brighton BN2 4GJ, UK; School of Immunology & Microbial Sciences, King’s College London, London SE1 9RT, UK

## Abstract

Organisation of the transcription cycle is facilitated by the reversible phosphorylation of the C-terminal domain of RNA Polymerase II (RNAPII-CTD) and its accessory factors. The PNUTS-PP1 protein phosphatase is crucial for mRNA synthesis and processing, yet the complete spectrum of its physiological targets in these processes remain elusive. Here, using quantitative phosphoproteomics, we discover that Cdk11, in addition to various spliceosomal and RNA processing factors, associates with PNUTS, and that disruption of PP1-binding results in hyperphosphorylation of Cdk11 at an evolutionarily conserved Serine residue, seven amino acid residues C-terminal to DFG residues in the activation loop. *In vitro* experiments reveal a role for Ser DFG+7 in modulating Cdk11 kinase activity towards RNAPII-CTD Ser5. Making use of a novel technique to conditionally disrupt PP1 binding, we show that PNUTS-PP1 normally serves to restrain Cdk11 phosphorylation *in vivo*. Mutational analysis shows that *cdk11* is not only essential for survival but also plays a widespread role in regulating normal mRNA expression and splicing. Notably, we find that a phosphomimetic mutation in *cdk11* exhibits distinct biological effects compared to loss of *cdk11* function, including defective processing of intronic small nucleolar RNAs, diminished intronic RNA Pol II velocity, and a decrease in intergenic transcription. These findings underscore physiologically significant roles of Cdk11 dephosphorylation by PNUTS-PP1 in the regulation of mRNA transcription and processing.

## Introduction

Protein phosphatase 1 (PP1) is one of the major serine/threonine protein phosphatases in eukaryotic cells and plays many essential physiological roles, including regulation of transcription and mRNA splicing [1–3]. Although it possesses pleiotropic functions, the catalytic subunit of PP1 interacts with more than 200 structurally and functionally diverse interacting proteins (PIPs) *in vivo* that define PP1’s specific roles at distinct subcellular locations [4–7]. Interaction with PP1 is largely mediated by a canonical PP1-binding motif common to most interacting proteins [8], such that each PP1-PIP complex defines a unique holoenzyme. The PP1 Nuclear Targeting Subunit (PNUTS) is one of the two major PIPs in the mammalian nucleus, and targets PP1 to RNA Polymerase II (RNAPII) at active sites on *Drosophila* chromosomes [9]. Profiles of chromatin-binding in mammalian cells indicate that PNUTS is predominantly located at relatively highly expressed genes including metabolic regulators [10], as well as histone clusters and small nucleolar RNA (snoRNA) genes [10, 11]. PNUTS is enriched downstream of the transcription start site (TSS) of many genes, peaking around the RNAPII pause site and at the first exon-intron boundaries, and is also enriched near transcription end sites (TES) [11]. PNUTS localisation at sites of active transcription is mediated in part via association with nascent RNA in mammals [11] and in *Drosophila* (our unpublished observations), which is consistent with its possession of multiple C-terminal RNA-binding motifs, and the finding that targeting to specific genes can be mediated by long noncoding RNAs (lncRNA) [12, 13].

Functional studies have revealed an important requirement for PNUTS-PP1 in regulating RNAPII-carboxy-terminal domain (CTD) Ser5 phosphorylation [9, 14], a modification linked to initiating and early-elongating RNAPII, 5’ mRNA capping, co-transcriptional splicing and RNAPII turnover [15, 16]. In *Drosophila*, elevated Ser5 phosphorylation resulting from loss of PNUTS function is accompanied by global changes in expression of many genes, particularly those that are highly expressed, leading to defects in developmental growth and larval lethality [9]. The specific roles of RNAPII-CTD Ser5 dephosphorylation by PNUTS-PP1 are not yet well understood, although it may help facilitate Pol II turnover during the resolution of transcription-replication conflicts in human cells [17]. Other studies have begun to pin-point specific functions for PNUTS in the transcription cycle. In mammalian cells, PNUTS/PP1 plays a role in a first exon termination checkpoint by stimulating transcription termination of cis-regulatory sequences during early elongation [18]. PNUTS also facilitates transcriptional termination of unstable non-coding RNAs (ncRNAs) by the Restrictor complex, comprised of ZC3H4, WDR82, and ARS2, working with Symplekin, to synergistically dampen extragenic transcription [19, 20]. Notably, during transcription termination, PNUTS/PP1 dephosphorylates Spt5 to slow down RNAPII elongation downstream of polyA sites to facilitate targeting of RNAPII by the Xrn2 terminator exonuclease [21]. PNUTS has also been implicated in co-transcriptional mRNA splicing [11, 12] by inhibiting PP1 activity towards the splicing factor Sf3b1 to stimulate spliceosome activity [11]. In these ways, PNUTS has emerged a key player in mRNA synthesis and processing, coupling transcription elongation and termination with surveillance of ncRNAs, mRNA splicing and 3’ mRNA processing by modulating the phosphorylation status of RNAPII and its associated factors.

The identity of the regulatory kinases that work in conjunction with PNUTS-PP1 to determine the phosphorylation state of their common targets *in vivo* is poorly understood. For instance, multiple cyclin-dependent kinases (Cdks) are known to regulate phosphorylation of RNAPII-CTD. Phosphorylation of Ser5 by Cyclin-dependent kinase 7 (Cdk7), part of the Transcription Factor IIH (TFIIH) complex, marks the early stages of transcription and plays a pivotal role in the transition from transcription initiation to elongation by recruiting various RNA processing factors to the transcription machinery [22]. Another significant RNAPII-CTD kinase is Cdk11, which has roles in transcription elongation, mRNA splicing [23-25] and 3’ mRNA processing [26-30]. Cdk11 regulates intron removal of pre-mRNAs, interacting with and phosphorylating splicing factors RNPS1 and 9G8 [23, 24, 31] and Sf3b1[27], which triggers spliceosome activation [27]. The association with 3’ processing factors, FLASH and TREX, have also revealed roles for Cdk11 in 3′ end cleavage of replication-dependent histone mRNAs and HIV RNAs, respectively [26, 28, 32]. However, despite their shared substrates, little is known about the functional interplay between Cdks and PNUTS during gene expression.

While serine/threonine protein phosphatases, such as PNUTS-PP1, have emerged as critical and specific regulators of gene expression and cell function, there remains a limited grasp of their physiologically relevant substrates, which are critical for assessing phosphatase activity and understanding the molecular mechanisms that drive distinct biological outcomes of the different holoenzymes. To address these obstacles, researchers have developed strategies such as substrate trapping [33], which successfully identified SPT5 and RNAPII as PNUTS-PP1 targets [21, 33]. In our efforts to uncover novel interactors and substrates of PNUTS-PP1, we have employed a different approach, leveraging quantitative (phospho)proteomics to interrogate the phosphorylation states of stably associated PNUTS-interacting proteins with and without PP1 binding. Implementing this approach here has led us to identify a conserved Ser residue in the activation loop of Cdk11 as a substrate. We further corroborate this finding using a conditional PNUTS allele designed to disrupt PP1 binding upon activation. *In vitro* and *in vivo* analyses of Cdk11 phospho-site mutations underscore the significance of Cdk11 dephosphorylation and point to multifaceted roles of activation loop (de)phosphorylation in transcriptional regulation.

## Results

### The PNUTS-interactome in *Drosophila* embryos is enriched in RNA processing factors

PNUTS has been proposed to act as a scaffold for the assembly of multi-protein complexes containing PP1 [34], but the broader interactome for metazoan PNUTS and the effect of PP1 on the phosphorylation state of interacting proteins had not been described. Interaction of PP1 with PNUTS is mediated by a canonical K/R-K-(S/T)-V/L-T-W motif. Mutation of this motif by substitution of Trp to Ala greatly reduces PP1-binding [9]. Therefore, to identify the *Drosophila* PNUTS interactome in the presence or absence of PP1-binding, we ectopically overexpressed and immunoprecipitated HA-tagged PNUTS^WT^ and PNUTS^W726A^ proteins from *Drosophila* embyros. To reduce the isolation of non-specific nucleic acid-binding proteins, we treated embryo lysates with a DNA/RNA endonuclease prior to immunoprecipitation. For quantitative comparison of (phospho)proteins associated with PNUTS^WT^ and PNUTS^W726A^, we employed an isotope dimethyl labelling strategy for our trypic peptides [35] (**Figure 1A**). Four sets of PNUTS^WT^ and PNUTS^W726A^ IPs were differentially dimethyl labelled, generating a mass increase of 28.03 and 32.06 Da for light and intermediate dimethyl labels, respectively. To determine the PNUTS interactome, an equimolar mix was then analysed by liquid chromatography tandem mass spectrometry (LC-MS/MS) and compared to unlabelled control IPs (from cells not expressing PNUTS-HA). Two independent approaches were then used to define high-confidence protein interactors based on either prevalence or abundance of identification in PNUTS IPs compared to control IPs (see Methods). After filtering, 221 and 153 candidate PNUTS interactors remained, respectively, and were analysed in STRING to visualise the PNUTS interactor network (**Figure 1B**). This identified orthologues of a core PNUTS-PP1 complex [34], including the three main PP1 isoforms in *Drosophila* (Flw/PP1β-9C, PP1-87B and PP1-96A [36]) and *Drosophila* Wdr82 [9]. We did not identify Tox4, which may play a tissue-specific function (our unpublished findings). We also did not identify RNAPII, consistent with our unpublished observations that its interaction with PNUTS is RNA-dependent.

**Figure 1:**
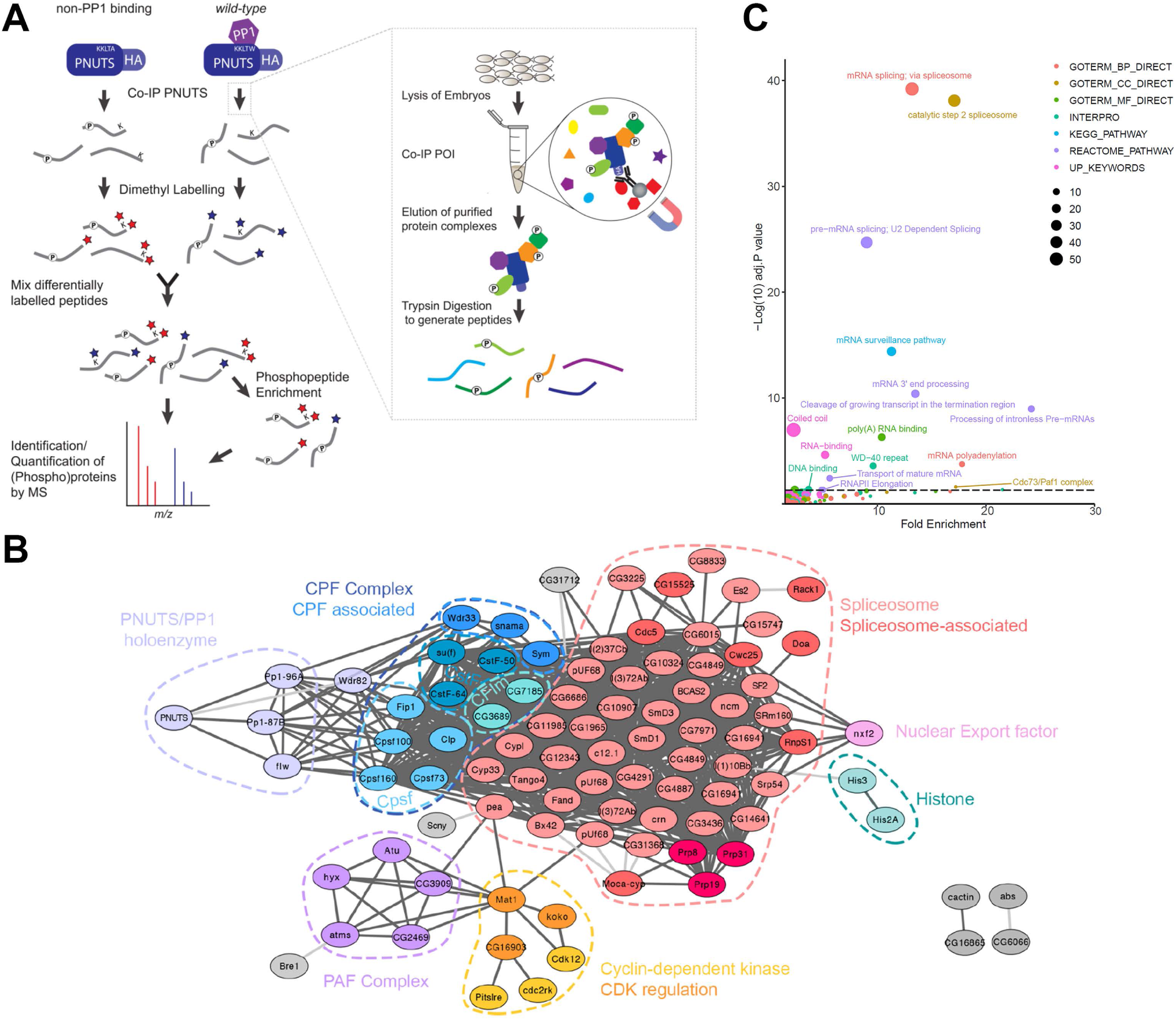
The Drosophila PNUTS-interactome. **A**, Illustration of the purification and labelling strategy to identify the PNUTS-interactome in *Drosophila* embryos expressing HA-tagged wildtype PNUTS (PNUTS^WT^) or a non-PP1 binding mutant (PNUTS^W726A^) PNUTS. Proteins were co-immunoprecipitated using magnetic beads conjugated to anti-HA antibodies (co-IP, see inset), and were eluted and digested with Trypsin. Resulting peptides were differentially dimethyl labelled and mixed prior to LC-MS/MS analysis; a proportion of the labelled peptide mixtures was enriched using TiO_2_ to enable quantification of phosphopeptides. Peptide signals from PNUTS^WT^ and PNUTS^W726A^ co-IPs were distinguished in LC-MS/MS by the difference in their mass. **B**, STRING network analysis of proteins identified in 2 or more PNUTS co-IPs with an average EmPAI score ≥2-fold higher than control co-IPs from a *w^1118^* strain. Edges connecting nodes represent either experimental or database evidence of high (interaction score >0.7, dark grey lines,) or medium (interaction score >0.4, light grey lines,) confidence in the STRING database. Group nodes are coloured according to functional groups/complexes. Nodes that did not fit into one of the annotated groups were coloured grey. **C**, Gene Ontology (GO) enrichment analysis of PNUTS interactors showing fold enrichment for significant GO categories after adjusting for multiple hypothesis testing using the Benjamini-Hochberg correction. The horizontal line represents an adjusted p value of 0.05. Points are sized according to the number of proteins within each enriched term and coloured according to the enrichment category.

In addition to identifying known PNUTS-interacting proteins, we found PNUTS precipitates were significantly enriched in RNA processing factors related to mRNA splicing (adj.p <0.001), 3’ processing (adj.p <0.001), polyadenylation (adj.p <0.001), mRNA transport (adj.p = 0.004) and RNAPII transcription elongation (adj.p =0.0046) (**Figure 1C**). This included representative components of almost all spliceosome subcomplexes [37], including all components of catalytic C complex spliceosome proteins (8/8 components). Precipitates were also enriched for Prp19/CDC5L proteins (6/17 members), and pre-catalytic/activated B/B* spliceosomal proteins (3/9 components). PNUTS also associated with all integral components of the polyadenylation factor complex (PAF1c)[38, 39]. We also found the majority of proteins comprising the core 3’ mRNA cleavage and polyadenylation factor (CPF) complex in our co-precipitates [40]. Also associated with PNUTS was a selection of cyclin dependent kinases (Cdks) linked to transcription and mRNA processing [23, 29, 41], including Cdk12, Pitslre/Cdk11 and cdc2rk/Cdk10, as well as Cdk-regulatory proteins including Mat1/MNAT1, koko/Cyclin Q, and CG16903/Cyclin L.

### Loss of binding to the major PP1 catalytic subunit isoform is the only significant change to the PNUTS interactome in PNUTS^W762A^ precipitates

Since IPs with PNUTS^WT^ and PNUTS^W726A^ were differentially dimethyl-labelled, we were able to directly compare the abundance of interacting proteins in PNUTS complexes in the presence or absence of an intact PP1-binding motif. Quantification of PNUTS interactors with respect to the amount of precipitated PNUTS protein showed that the only significant change was PP1-87B, the major PP1 isoform in *Drosophila* [42], which, was 12-fold more abundant in PNUTS^WT^ pull-downs than in those with PNUTS^W726A^ (*P* adj.=0.03 with Benjamini-Hochberg correction). Binding to Flw/PP1β9C [36] was also reduced, although below the level of significance (**Figure 2A**). Whilst not statistically significant, the majority of quantified PNUTS interactors were more abundant for PNUTS^W726A^ (low PP1 binding), indicating that PP1 is not the primary driver for binding of PNUTS to interacting partners in embryo extracts.

**Figure 2:**
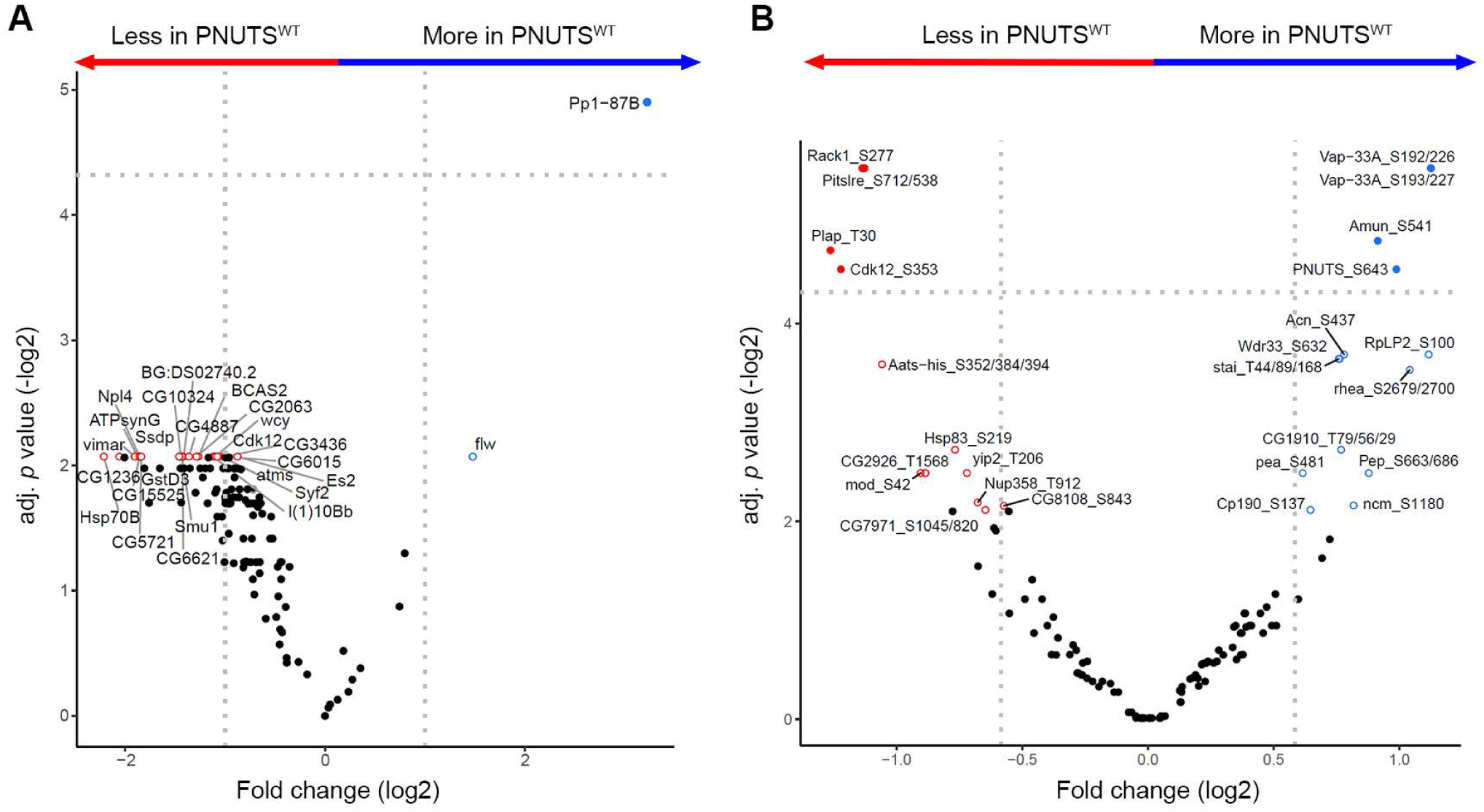
Effect of PP1-binding on the PNUTS-(phospho)interactome. Volcano plots showing the change in relative abundance of (**A**) PNUTS-interacting proteins and (**B**) phosphorylation of PNUTS-interacting proteins from PNUTS^WT^ and PNUTS^W726A^ precipitates, determined by quantitation of differentially dimethyl-labelled peptides. Log2 fold change indicates the difference between wildtype and non-PP1 binding conditions, where a positive fold change indicates that there was more protein (**A**) or phosphoprotein (**B**) present in co-IPs with PNUTS^WT^ compared to PNUTS^W726A^ (Blue arrow); a negative fold change indicates the converse relationship with more protein present in co-IPs with PNUTS^W726A^ compared to PNUTS^WT^ (Red arrow). Vertical dotted lines represent a 1.5-fold change in either direction. The horizontal lines represent an adjusted *p* value of 0.05 (Bayesian analysis with Benjamini-Hochberg *p* value correction). Filled blue and red dots represent points significantly (adj.*p*≤0.05) up or down in the PNUTS^WT^ respectively. Dots with red or blue outlines represent points with a significant *p* value (*p*≤0.05) in either direction prior to adjustment. Black points represent proteins whose abundance or phosphorylation status was not significantly altered in the presence/absence of PP1 (*p*>0.05).

### Analysis of the PNUTS-PP1 phospho-interactome reveals potential phosphatase substrates

Next, we sought to better understand the role of PP1 in regulating the PNUTS phosphoprotein-interactome. TiO2 enrichment of stable isotope dimethyl-labelled phospho-peptides (Figure 1A) was performed on PNUTS^wt^ and PNUTS^W726A^ IPs. To account for differences in protein abundance between PNUTS^WT^ and PNUTS^W726A^ precipitates, differences in phosphopeptides levels were normalised to fold change at the protein level. Using this approach, we were able to accurately quantify differences in phosphopeptide abundances in co-precipitates of PNUTS^WT^ and PNUTS^W726A^ (**Figure 2B**). Phosphorylated proteins that could not be quantified from the unenriched data in more than one sample were excluded from this analysis, to remove false positive identifications and allow for statistical comparisons.

Although the complete structure of the PNUTS:PP1 holoenzyme is yet to be determined, PNUTS’ high affinity PP1-binding sites are known to block one of PP1’s substrate binding grooves, effectively inhibiting dephosphorylation of some substrates, such as retinoblastoma protein (Rb), while leaving the active site accessible [43]. Another PP1 cofactor, Phactr1, docks with PP1 in a similar way to PNUTS, also occluding its C-terminal groove [44]. Notably, Phactr1 also creates a new surface pocket adjacent to the PP1 catalytic site [44], explaining how PIPs may facilitate dephosphorylation of specific substrates via composite PIP:PP1 substrate binding sites. Consequently, we aimed to differentiate between putative substrates of the PNUTS:PP1 holoenzyme (anticipated to be hyperphosphorylated with PNUTS^W726A^) and proteins belonging to PNUTS-associated complexes whose dephosphorylation by PP1 is normally inhibited by PNUTS (anticipated to be hyperphosphorylated with PNUTS^WT^). Consistent with this, there was partial enrichment of phosphopeptides containing a consensus recognition sequence for the PP1 catalytic subunit in proteins that exhibited enhanced phosphorylation with expression of PNUTS^WT^, and this may be indicative of PNUTS-dependent inhibition of PP1 phosphatase activity [45]. Amongst these proteins, we observed a 1.7-fold increase (P<0.01) in phosphorylation of the splicing factor Acinus at Ser 437 in PNUTS^WT^ compared to PNUTS^W726A^ IPs. Acinus pSer 437 was previously identified as a putative PP1 substrate in a screen for genetic modifiers of *acinus* overexpression [46]. Notably, we did not identify differential phosphorylation of Sf3b1. The N-terminus of human Sf3b1 is specifically phosphorylated at multiple residues, including Ser 129 and Thr 313, during human spliceosome activation [47-49]. It has been proposed that PNUTS inhibits Sf3b1 dephosphorylation by the NIPP1-PP1 holoenzyme, by binding PP1 in place of NIPP1 [11, 50]. Although we detected phosphorylated *Drosophila* Sf3b1 peptides (pSer 137, equivalent to Ser 129 in human Sf3b1) in our IPs, indicating that PNUTS may localise to activated spliceosomes, we did not observe any significant difference in Sf3b1-Ser137 phosphorylation between PNUTS^WT^ and PNUTS^W726A^ precipitates (P=0.34). We detected several hyperphosphorylated proteins in PNUTS^W726A^ precipitates. One of the most prominent changes (2.2-fold increase, P<0.001) in phosphorylation in PNUTS^W762A^ compared to PNUTS^WT^ precipitates was observed for a highly conserved Ser residue located in the activation loop of Cdk11/Pitslre (Ser 538/712 in *Drosophila* Cdk11 p89/p109 isoforms) [51].

### PNUTS-PP1 maintains Pitslre/Cdk11 in a dephosphorylated state *in vivo*

To confirm the role of PNUTS-PP1 in regulating Cdk11 phosphorylation, we examined the effect of manipulating PNUTS-bound PP1 activity *in vivo* in a specific and timely manner. To do this, we devised a genetic approach that we refer to as InDisruPT (Inducible Disruption of Phosphatase Targeting), which makes use of a conditional transgenic allele of PNUTS capable of being converted irreversibly from PP1-binding (PNUTS^WT^) to non-binding (PNUTS^W762A^). This allele contains a duplicated 3’ exon carrying the W762A mutation, which greatly reduces binding to PP1 ([9], **Figure 2A**). The duplicated exon is only expressed after FRT/FLP-induced recombination, which irreversibly replaces the wildtype exon, tagged with GFP, with the mutant version, tagged with mCherry (PNUTS^WT^GFP-PNUTS^W726A^mCh), allowing the conversion to be monitored post-translationally (**Figure 3A**). To control for the function of GFP verses mCherry-tagged PNUTS, an analogous switchable allele was made that converts PNUTS^WT^-GFP to PNUTS^WT^-mCherry (PNUTS^WT^GFP-PNUTS^WT^mCh), such that the expressed protein only differs by the C-terminal fluorescent tag.

**Figure 3:**
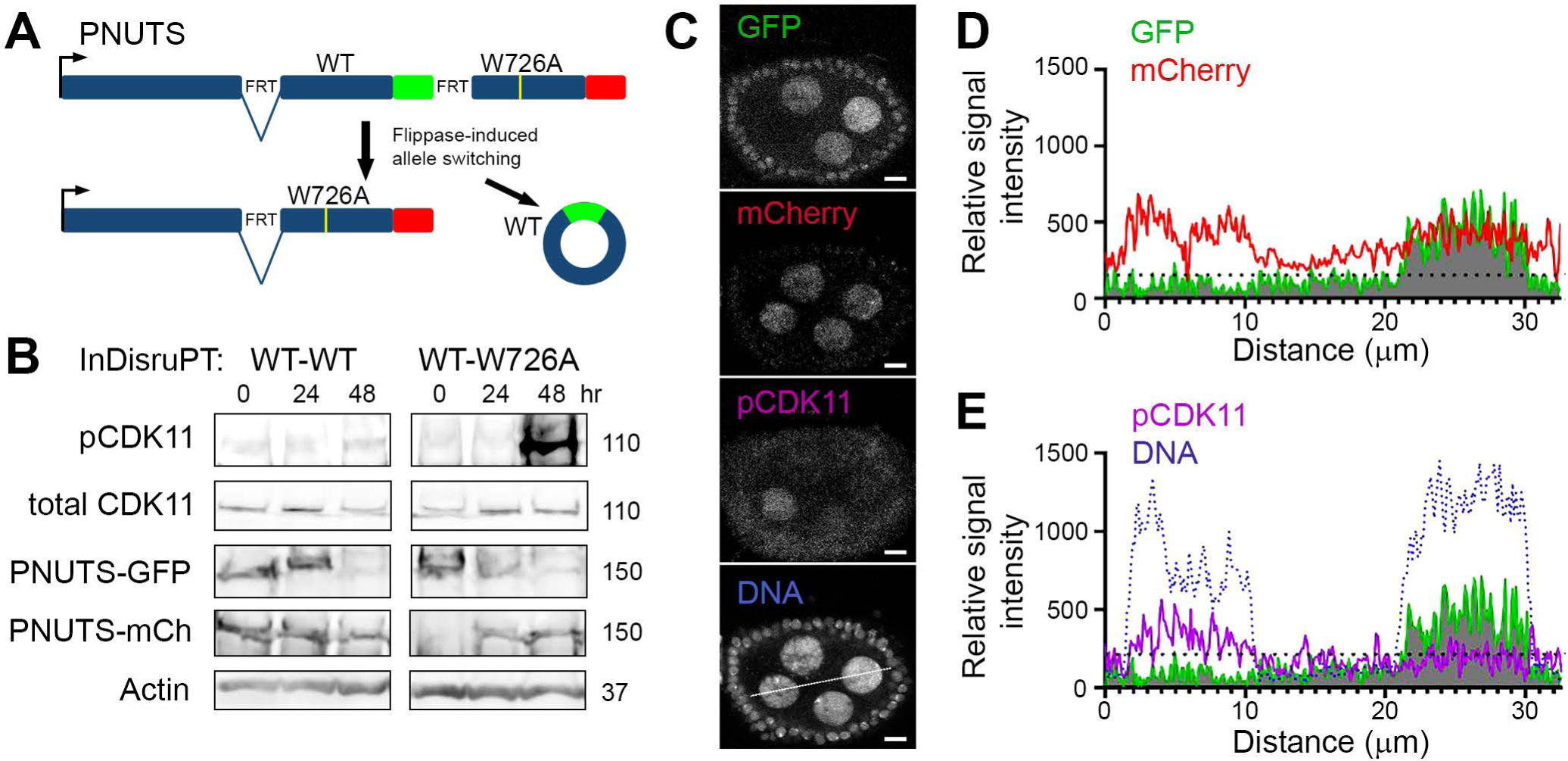
PNUTS-PP1 is required to restrict cdk11-S712 phosphorylation in vivo. **A**, Strategy for Inducible Disruption of Phosphatase Targeting (InDisruPT). Structure of a genomic transgene is depicted, before and after FLP-mediated recombination at FRT sites flanking a 3’ exon encoding the C-terminus of PNUTS. Shown are the wildtype 3’ exon, with GFP coding sequence in green, and non-PP1 binding variant, with mCherry in red. Recombination results in production of extragenic wildtype circular DNA, which is not maintained, and reconstruction of the PNUTS transcription unit carrying the alternative 3’ exon, which harbours the W726A mutation. **B**, Western blots showing that inducible disruption of PP1 binding to PNUTS results in Cdk11 Ser 712 phosphorylation *in vivo*. Adult flies carrying InDisrupt *PNUTS* transgenes, were induced with hsFLP and blotted with GFP, RFP, Cdk11-pSer 712, total Cdk11, and Actin antibodies either 0, 24, or 48hr after induction. **C**, Confocal images of early-stage egg chamber showing switching of PNUTS^WT^GFP: PNUTS^W726A^mCh specifically in germline cells using ovoFLP, and the effect on Cdk11-pSer712 staining. Switching is incomplete in some cells, as shown with a high GFP:RFP ratio. Scale bars, 50 µm. Line indicates position of line scan taken through the image for quantitation. **D**, Plots of line scan signal intensities for pCdk11, RFP, GFP and DNA (TO-PRO-3 dye) staining, showing effect of switching PNUTS^WT^GFP to PNUTS^W726A^mCh on Cdk11-pSer712 in nurse cell nuclei.

To test whether loss of PNUTS-PP1 dephosphorylates Cdk11 *in vivo*, we used InDisRuPT to switch PNUTS^WT^GFP-PNUTS^W726A^mCh in a genetic background lacking all endogenous *PNUTS* coding sequences (*PNUTS^13B^*/*PNUTS^13B^*, [9]). Partial turnover of PNUTS^WT^GFP and *de novo* expression of PNUTS^W726A^mCh was evident by GFP and RFP immunoblotting of extracts at 24 hr after FLP induction, and that this transition was complete by around 48 hr. Similar dynamics were evident for a control strain lacking the W726A mutation (PNUTS^WT^GFP-PNUTS^WT^mCh), although there was some leaky PNUTS^WT^mCh expression at an earlier stage (**Figure 3B**). Blotting with a custom-made antibody revealed a concomitant 5-fold increase (n=2) in the phosphorylation of the Cdk11 long isoform (p109) at Ser 712 in PNUTS^WT^GFP-PNUTS^W726A^mCh compared to PNUTS^WT^GFP-PNUTS^WT^mCh extracts in a *PNUTS^13B^* background. Total Cdk11 levels were relatively unaffected (**Figure 3B**). To confirm elevated Cdk11 phosphorylation following reduction of PNUTS/PP1 binding, we induced allelic switching of PNUTS^WT^GFP-PNUTS^W726A^mCh in ovarian germline cells and subjected dissected egg chambers to immunostaining (**Figure 3C**). Line scans through confocal images showed that fully switched PNUTS^W726A^ cells had higher pSer 712 Cdk11 levels compared to partially switched PNUTS^WT^GFP-PNUTS^W726A^mCh cells where both GFP and RFP signals were visible (Compare **Figure 3D & E**).

### Cdk11 Ser712 directly modulates Cdk11 activity towards RNAPII *in vitro*

Phosphorylation of a conserved Thr residue in the activation loop of cyclin-dependent kinases is known to be critical for kinase activation [51-54]. However, the regulatory role of the upstream Serine residue (+7 residues C-terminal to the canonical DFG motif), is poorly studied, except in human Cdk7, where phosphorylation of the analogous residue (S164) by CK1 or CK2 inactivates Cdk7 and impairs transcriptional activity of TFIIH [52, 53]. The DFG+7 Ser is highly conserved in Cdk11 orthologues, although the *S.pombe* enzyme contains a negatively charged Glu residue at this position (**Figure 4A**). To investigate the importance of Ser 712 phosphorylation on Cdk11 activity, we purified recombinant full-length N-terminally His-tagged *Drosophila* Cdk11:Cyclin L enzymes from insect Sf9 cells and probed for kinase activity towards the physiological substrates RNAPII-CTD and Sf3b1 [26-28, 30]. Recombinant RNAPII-CTD was incubated with either Cdk11^wt^ or a kinase dead variant (Cdk11^KD^, D703A) and immunoblotted with RNAPII-CTD antibodies that are specific for phospho-Ser2, 5, 7, Thr4, using total RNAPII-CTD as a control [55]. Cdk11^wt^/Cyclin L, specifically phosphorylated RNAPII-CTD Ser5 and none of the other Ser/Thr residues in the CTD repeat (**Figure 4B**). Importantly, Cdk11^KD^/Cyclin L did not phosphorylate RNAPII, confirming on-target kinase activity of Cdk11^wt^/Cyclin L. In contrast, we did not detect an increase in Thr 313 phosphorylation for recombinant human Sf3b1 incubated with Cdk11^wt^/CyclinL, or observe changes in *Drosophila* Sf3b1-pS137 signals upon *cdk11* knockdown in *Drosophila* extracts.

**Figure 4:**
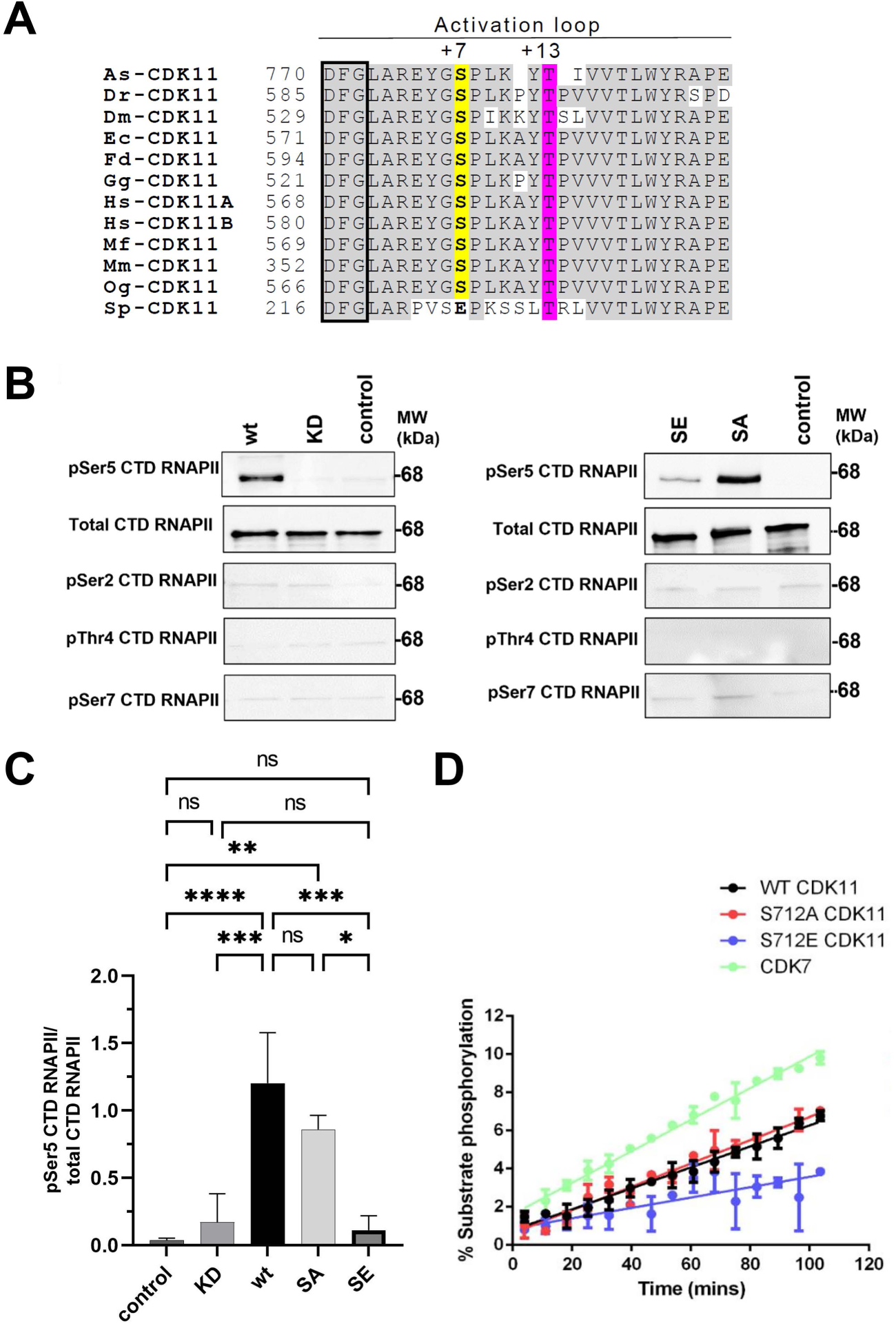
Modification of Serine DFG+7 alters Cdk11 kinase activity towards RNAPII-CTD. **A**, Multiple sequence alignment of Cdk11 proteins showing evolutionary conserved features, including DFG motif (boxed), Serine DFG+7 (yellow highlight) and the activating Threonine at DFG+13 (magenta highlight). As, *Anopheles sinensis* (mosquito); Dr, *Danio rerio* (zebrafish); Dm, *Drosophila melanogaster* (fly); Ec, *Equus caballus* (horse); Fd, *Fukomys damarensis* (rat): *Gallus gallus* (chicken); Hs, *Homo sapiens* (human); Mf, *Marmota flaviventris* (marmot); Mm, *Mus musculus* (mouse); Og, *Otolemur garnetti*i (Galago); Sp, *Schizosaccharomyces pombe* (fission yeast). **B**, Effect of Cdk11 DFG+7 amino-acid mutations on Cdk11/Cyclin L activity towards recombinant RNAPII-CTD. RNAPII-CTD was incubated with or without recombinant WT or mutant variants of Cdk11/Cyclin L, before immunoblotting with RNAPII-CTD antibodies specific for pSer2, 5, 7 or pThr4 or total RNAPII-CTD. Cdk11^WT^/Cyclin L and Cdk11^S712A^/Cyclin L showed activity towards RNAPII-CTD Ser5, whereas Cdk11^S712E^/Cyclin L and Cdk11^KD^/Cyclin L showed reduced and no activity, respectively. **C**, Quantification of pSer5 RNAPII-CTD band intensities, normalised to total RNAPII-CTD (n=3). Data shown are mean ±SEM (n=3). **, p=<0.01, one-way Anova. **D**, Time-dependent phosphorylation of CTD peptide 5-FAM-(YSPTSPS)_3_-CONH_2_, by CDK7/Cyclin H (∼0.5μg, positive control) and Cdk11/Cyclin L1 variants (∼1μg enzyme). Mean ±SD, n=2.

Next, we sought to examine the effects of Ser712 phosphorylation on Cdk11 kinase activity towards RNAPII using non-phosphorylatable (Cdk11-Ser712Ala) or putative phospho-mimetic (Cdk11-Ser712Glu) recombinant enzymes. Notably, there was no significant difference in RNAPII-CTD Ser5 kinase signal when incubated with Cdk11^S712E^/Cyclin L or inactive Cdk11^KD^/Cyclin L (**Figure 4C**), whereas Cdk11^S712A^/Cyclin L retained 71.5±6.1% of the activity of Cdk11^wt^/Cyclin L. The catalytic activities of our Cdk11 variants towards a fluorescently-labelled substrate peptide derived from the RNAPII-CTD (5-FAM-(YSPTSPS)_3_-CONH_2_) was monitored kinetically in real-time using a microfluidic kinase assay system [56], using active CDK7 as a positive control. These assays confirmed depleted activity of Cdk11^S712E^/Cyclin L towards the Ser5 containing peptide, relative to WT and S712A variants (**Figure 4C**). Immunoblot analysis also confirmed Cdk11/Cyclin L specific phosphorylation of Ser5 in the RNAPII-CTD containing peptide.

### *In vivo* requirement for *cdk11* and cdk11^S538/S712^

To determine the *in vivo* role of *cdk11*, we generated a loss-of-function allele of *cdk11* by CRISPR-Cas9 mediated insertion of a dsRed-marked *PBac* element to disrupt the *cdk11* coding sequence (*cdk11^P^*)(**Figure 5A**). *cdk11^P^* homozygous flies were lethal as 1^st^ instar larvae, most likely once maternally provided *cdk11* became depleted (**Figure 5B**). Homozygous lethality of *cdk11^P^* animals was rescued to adulthood by complementation with a transgenic copy of genomic *cdk11*, confirming lethality was not due to off-target effects (**Figure 5C**). To determine the requirement for Cdk11 Ser 538/712 during development, we engineered phospho-mimetic (SE) or non-phosphorylatable (SA) mutations into our cdk11 genomic rescue construct and examined the ability of transgenic copies of these constructs, inserted at the same landing site as the wildtype transgene, to rescue the lethality of *cdk11^P^*/*cdk11^P^* larvae. Interestingly, genomic rescue was abrogated with the cdk11 ^S712E^ transgene compared to cdk11^WT^ and cdk11^S712A^ transgenes, which fully rescued larval lethality (**Figure 5C**).

**Figure 5:**
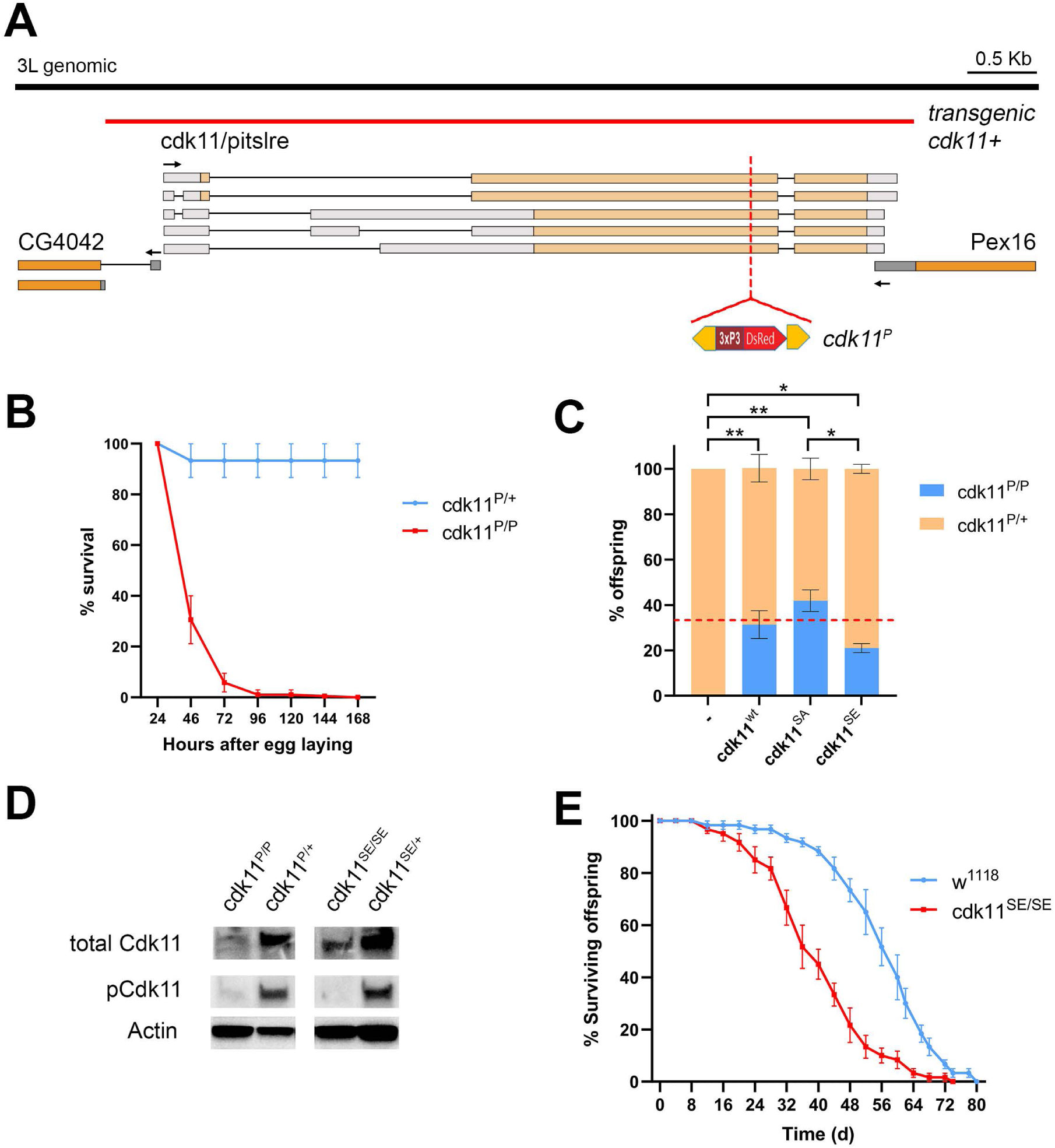
In vivo analysis of Cdk11 function in developing *Drosophila* larvae. **A**, Genomic region of *Drosophila cdk11* showing exon-intron structure for *cdk11* and spatial relationship to flanking genes on chromosome 3L. Orange shading represents coding regions, grey boxes represent untranslated regions. *cdk11^P^* contains an 3xPS>DsRed marked *PBac* element inserted into the coding sequence of *cdk11* by CRISPR-Cas9 mediated gene targeting. Removal of PBac generated seamless mutation of S538/712 (*cdk11^SE^*) due to site-directed mutations present in the homology arm. Also shown is the position of wild type genomic fragment containing the *cdk11* transcription unit, which were used to make transgenic lines (cdk11^wt^ and site-directed mutants cdk11^SA^ and cdk11^SE^). **B**, Graph showing percentage of hetero- and homozygous *cdk11^P^*animals surviving over time (days after egg laying). More than 90% of *cdk11^P^* died by 72 hr. **C**, Graph showing ability of cdk11 genomic transgenes to rescue lethality caused by *cdk11* loss-of-function. Shown are the proportion of offspring from genetic crosses that were either heterozygous (orange data points) or homozygous (blue data points) for *cdk11^P^*. Mean ±SEM of three biological repeats is shown. The expected number homozygous *cdk11^P^*animals upon complete rescue by *cdk11* transgenes (33%) is indicated with a red dashed line. Both *cdk11^wt^* and non-phosphorylatable *cdk11^SA^*transgenes fully rescued *cdk11^P^* lethality, whereas only partial rescue was observed with phospho-mimetic transgenic *cdk11^SE^*. *, p<0.05, ** p<0.001 one-way Anova. **D**, Western blots of protein lysates from homozygous or heterozygous *cdk11^P^* and *cdk11^SE^* mutant larvae, showing loss of Cdk11 protein in *cdk11^P^* extracts. Actin immunoblots show loading control. pCDK11 blots show lack of signal in either *cdk11^P^* and *cdk11^SE^* homozygous extracts, further confirming specificity of this antibody for phospho-Cdk11. **E**, Plot showing survival of *cdk11^SE^* homozygous males (median survival 39.4 days) compared to *w^1118^* control (median survival 58.4 days).

These data support a physiological role for Ser 538/712 phosphorylation in regulating Cdk11 function. To provide further evidence of this, we used the CRISPR-Cas9 homology arms of our *PBac* insertion line to introduce a seamless Ser-Glu mutation (*cdk11^SE^*) after PBac excision. Immunoblotting confirmed loss of Cdk11 expression in the parental line (*cdk11^P^*) and re-expression of Cdk11 in the excision line (cdk11^SE^) (**Figure 5D**). Although homozygous *cdk11^SE^* animals were able to survive to adulthood, they displayed reduced longevity compared to *w^1118^*controls (**Figure 5E**).

### *cdk11* loss of function and phosphomimetic mutations induce both common and distinct changes in mRNA abundance

To better understand the biological effect of *cdk11* mutations, we used RNA-Seq to analyse the transcriptome of wild type and mutant homozygous larvae. For this, we processed total mRNA to capture information about transcripts irrespective of their polyadenylation status. Unsupervised clustering of normalised gene expression data based on sample dissimilarity showed replicates for each genotype were highly related to each other and indicated divergent transcriptional responses between *cdk11^P^, cdk11^SE^* and their developmentally matched (*w^1118^*) controls (**Figure 6A**). Differential gene expression (DGE) analysis revealed 2,772 and 2,910 genes in *cdk11^P^*and *cdk11^SE^* larvae, respectively, were significantly affected (padj<=0.05) by more than 1.5 fold, indicating a shift in transcription state in animals at the point at which they were analysed (**Figure 6B**). We observed only partial overlap in differentially expressed genes and enriched gene ontology terms between the two mutant alleles, although underexpressed genes in both *cdk11^P^* and *cdk11^SE^* were enriched in metabolic genes (**Figure 6C**). A notable difference between *cdk11* alleles was that 91% of snoRNAs that we could detect in controls (n=105) were strongly underexpressed in *cdk11^SE^* (on average 2.38 ±0.12 log2-fold reduced compared to controls), whereas the mean snoRNA expression level was unaffected in *cdk11^P^* homozygotes (0.29 ±0.04 log2-fold overexpressed compared to control)(**Figure 6D**, bottom). The majority of snoRNA genes are generated through alternative splicing of host genes in which they reside. Strikingly, whilst the level of intronic snoRNAs was significantly affected in *cdk11^SE^*animals, expression of the respective host genes was not (**Figure 6D**, top), indicating that host gene and snoRNA expression was effectively uncoupled in this strain.

**Figure 6:**
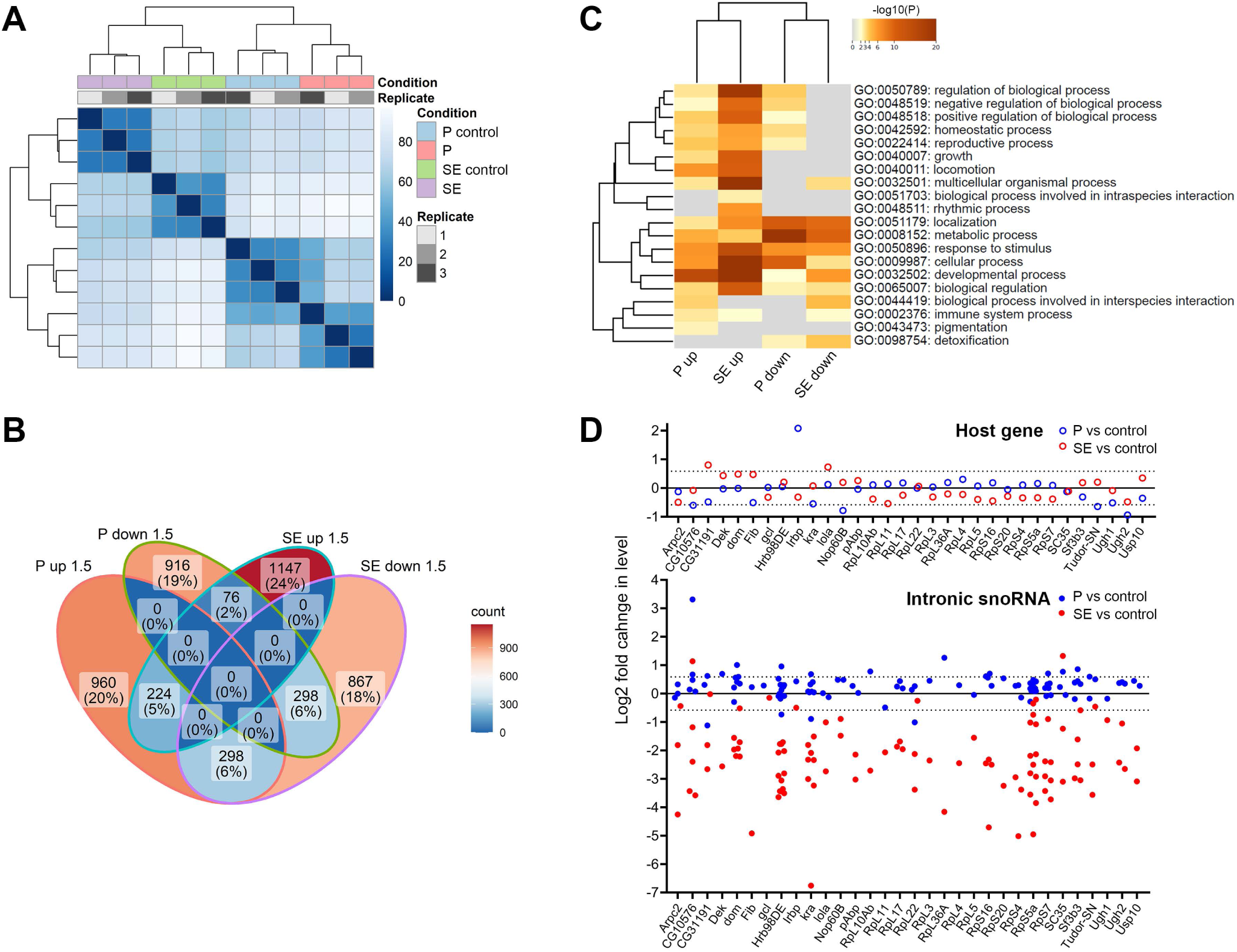
Transcriptomics reveals differential gene expression in *cdk11* mutants. Total RNA from *cdk11^P^* and *cdk11^P^* homozygous larvae and developmentally-matched controls were analysed by RNA-Seq. **A**, Unsupervised hierarchical clustering of samples based on their DESeq2 normalized gene-level counts shows the degree of similarity between samples. The heat map displays inter-sample Euclidean distances, where darker blue colours indicate closer similarity. **B**, Venn diagram showing the overlap between the genes significantly differentially expressed in *cdk11^P^* or *cdk11^P^*(more than 1.5 fold increased or decreased compared to controls, (p adj<=0.05) showing common and divergent responses. Numbers of genes in each intersect are indicated, along with the percentage of genes represented from each genotype. **C**, Gene Ontology enrichment amongst significantly overexpressed or underexpressed genes in *cdk11^P^* or *cdk11^P^* generated using Metascape. The scale bar indicates the negative log(10) p-values for enrichment. Only significant categories are shown, and grey coloured cells indicate non-significance. Metabolic process and Response to stimuli are commonly enriched categories. **D**, Plots of log(2) fold change in expression of intronic snoRNAs (bottom) and their respective host genes (top) in *cdk11^P^*(blue data points) or *cdk11^SE^* (red data points) larvae.

### cdk11 mutations display distinct intronic and intergenic transcriptional signatures

Since Cdk11 is a known splicing regulator, we examined intron retention by measuring the percent spliced-in (PSI) exons [57, 58] in transcripts from mutant animals. Notably, there was more intron retention in *cdk11^P^*mutant larvae (n=427 events), than in *cdk11^SE^* (n=22 events), when compared to controls (**Figure 7A-B**). To examine this further, we examined the saw-tooth pattern of intronic RNA-Seq read coverage, which is indicative of Pol II speed and co-transcriptional splicing. When RNAPII operates at high speeds there are fewer instances of nascent transcripts being interrupted within introns at the time of cell lysis, resulting in a less steep decline in read coverage from the 5’ to 3’ end of an intron. *cdk11^P^* had only a modest effect on the estimated RNAPII speed by this measure (1.1-fold reduction, n=403 introns sampled), whereas in *cdk11^SE^* animals, intronic RNAPII speed was reduced on average 2.4-fold (CI ±0.2) (n=415 introns sampled) (**Figure 7C**). Since PNUTS has been linked to RNAPII speed and transcription termination [19, 21, 59, 60], we also measured the percentage of sequence reads mapping to intergenic bases in the *Drosophila* reference genome. We found that the mean percentage of intergenic bases was elevated 3.07-fold (P=0.018) in cdk11^P^ and reduced 1.48-fold (p=0.054) in *cdk11^SE^* compared to matched controls (**Figure 7D**). This prompted us to investigate the source of intergenic sequences in *cdk11^P^* further by analysing the global level of readthrough transcription in each sample using ARTdeco, which measures the ratio of read-in (1-15 kb upstream of reference genes) and readthrough (4–15 kb in length ) transcription for the top 1000 expressed genes [61]. We found a significant 1.28-fold increase in the read-in summary statistic in *cdk11^P^* homozygotes compared to matched controls (P=0.039), accompanied by a 2.29-fold increase in the summary statistic characterising downstream-of-gene transcription (P=0.022) (**Figure 7E, 7G**). In contrast, *cdk11^SE^* showed significant 1.65-fold (P=0.023) and 2.02-fold (P=0.0084) reductions in transcriptional read-in and readthrough, respectively, compared to controls (**Figure 7E, 7F**). Together, these data point to the potential for multiple roles for *Drosophila cdk11* in the control of mRNA synthesis and processing, and a distinct requirement for Cdk11 dephosphorylation in these processes.

**Figure 7:**
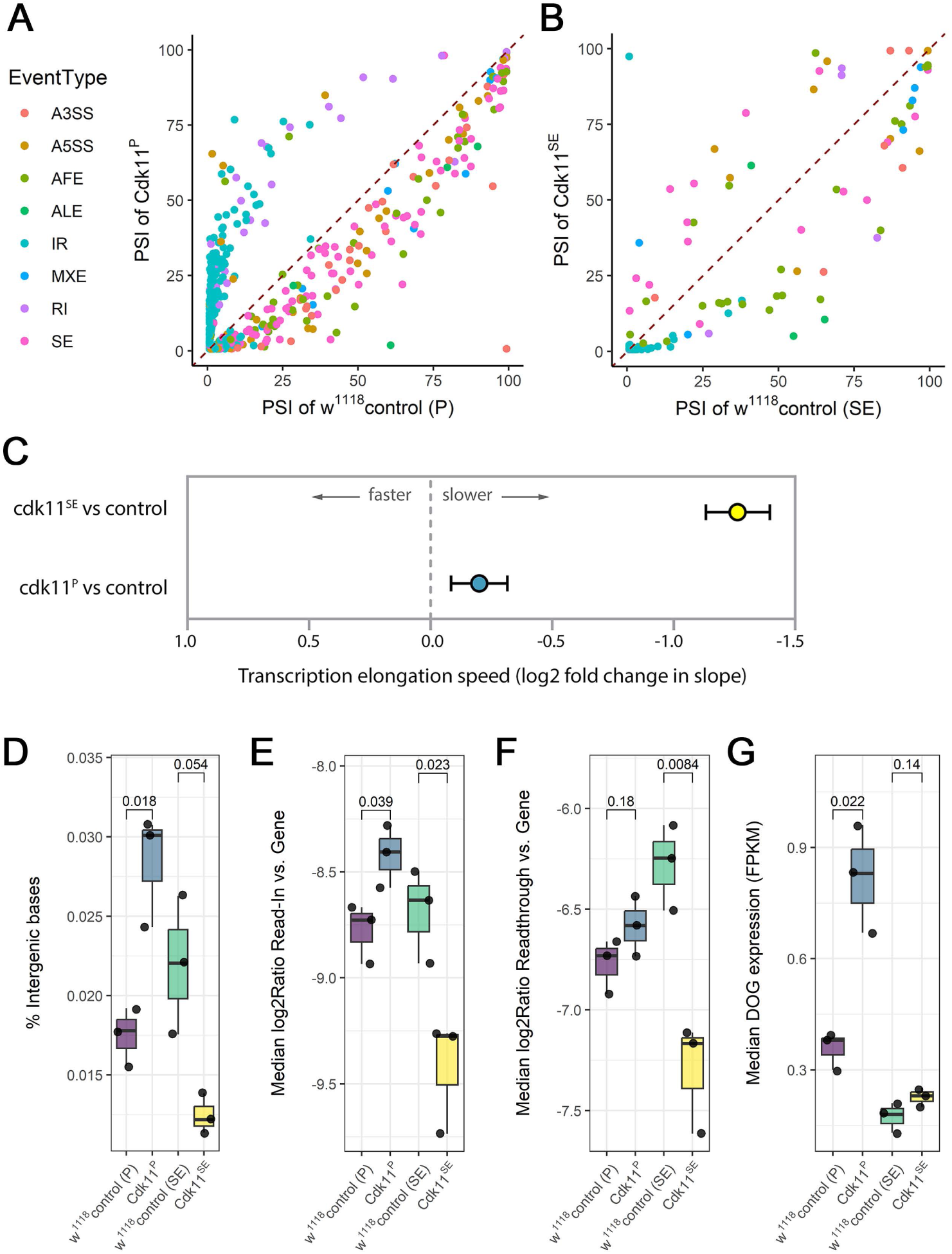
*cdk11* mutants differentially affect RNAPII transcription rate, RNA processing and termination. **A,B**, Percent spliced in index (PSI) plots showing significant differences (*padj*<=0.05) in splicing events identified between the control and *cdk11* mutant strains. Colour key indicates different categories of splicing events that were examined using NxtIRF: intron retention (IR), mutually exclusive exons (MXE), skipped exons (SE), alternate first exon (AFE), alternate last exon (ALE), alternate 5’-splice site (A5SS) and alternate 3’-splice site (A3SS). Intron retention events (IR, cyan colour) are increased in *cdk11^P^*compared to its matched control (**A**). In contrast, there were fewer intron retention events in *cdk11^SE^* animals compared to the control (**B**). **C**, Plot of mean difference in RNAPII elongation speed in *cdk11^P^* and *cdk11^SE^* compared to the respective controls. The log-2 fold change in mean coverage between mutant and control samples is shown, oriented so that negative values represent more negative slopes and slower estimated transcription in the mutants. **D,** *cdk11^P^* animals show an increase in aberrant intergenic transcription, as measured by the percentage of RNA-Seq reads mapping to intergenic bases (% intergenic bases, PICARD). T-test p-values are indicated above each mutant-control group comparison (n=3). **E-G**, *cdk11* mutants differentially modulate transcriptional read-in and readthrough compared to control animals. **E**, Boxplot showing the median log(2) ratio of read-in expression (1-15 kb upstream of genes) to gene expression. **F**, Boxplot showing the median log(2) ratio of read-through expression (downstream, 4-15 kb in length) to gene expression. **G**, Boxplot of median downstream of gene expression (FPKM). In each panel, the results of T-tests between matched mutant-control pair groups are shown (n=3 per group).

## Discussion

### The PNUTS (phospho)interactome

Recent studies point to multiple roles for PNUTS-PP1 during mRNA expression, from RNAPII elongation and termination to RNA splicing. To better understand the engagement of PNUTS and PNUTS-bound PP1 with these core processes, we sought to identify the PNUTS interactome and phospho-interactome in *Drosophila*. The isolation of PNUTS-associated factors belonging to active spliceosome, Paf1 and CPF complexes, is consistent with PNUTS-PP1’s multifunctional roles during the mRNA transcription cycle. Binding to these factors in the absence of RNA/DNA rules out non-specific interactions via nascent RNA, indicating instead that PNUTS stably associates with these complexes. CPF components have repeatedly been found to co-purify with the spliceosome and play roles in alternative splicing [37, 62, 63]. Therefore, understanding the intricacies of PNUTS interactions with these components will demand further investigation to elucidate which components are directly responsible for these connections.

Identifying physiological substrates of protein phosphatases is an important challenge. Although the *in vitro* substrate preference of PP1 catalytic subunit has recently been described, *in vivo* PP1 exists as more than 200 distinct enzymes defined by their interaction partner(s) [4-7]. Each PP1 holoenzyme possesses the ability to confer specificity to the PP1 catalytic subunit by modulating substrate-binding sites, with positive or negative effects on substrate binding, or by enriching the phosphatase at specific subcellular sites [4, 5]. To address this issue, we focused on potential substrates that stably associate with PNUTS in the presence of absence of PP1 binding. One limitation of this approach is that it potentially overlooks substrates with transient associations, or those whose interactions are contingent on phosphorylation states. Also, some substrates may depend on RNA for their association with PNUTS-PP1, such as RNAPII. Our analysis identified proteins whose phosphorylation state increased or decreased in the absence of PP1-binding to PNUTS, consistent with PNUTS acting as both an inhibitor and substrate specifier, depending on the protein substrate.

### Emergence of the PNUTS-PP1/CDK11 regulatory module determining RNAPII Ser5 phosphorylation

Foremost amongst the targets for dephosphorylation by PNUTS-PP1 that we identified was Cdk11. Modelling the effect of phosphorylation *in vitro*, we found that non-phosphorylatable Cdk11/Cyclin-L (*cdk11^S712A^*) possesses more RNAPII-CTD-Ser5 kinase activity than a phosphomimetic allele (*cdk11^S712E^*). Strikingly, this suggests that PNUTS-PP1 activity has the potential to both reduce RNAPII phosphorylation through its own RNAPII-directed phosphatase activity or stimulate increased phosphorylation indirectly by dephosphorylating Cdk11. Self-limiting behaviour of this type may be triggered by a timing mechanism or in response to stimuli, such as the recruitment of Cdk11 to RNAPII. Whether net RNAPII-CTD-Ser5 is elevated or reduced by PNUTS-PP1 may also depend on recruitment of antagonistic kinases with the potential to switch off Cdk11 through phosphorylation of Ser DFG+7, thereby promoting net RNAPII-CTD-Ser5 dephosphorylation by PNUTS-PP1. Deciphering such networks will require careful analysis of the effects of acutely perturbing kinase and phosphatase activities in an appropriately tractable system.

Intriguingly, sequence comparisons of Cdk11 and PNUTS orthologues point to the possible evolutionary origins of RNAPII modulation by Cdk11/PNUTS-PP1. Budding yeast does not possess a Cdk11 homologue, while *S.pombe* Cdk11 possesses a Glu residue at the DFG+7 position (Fig.4A, as in our *cdk11^S712E^* allele) and has been reported to lack RNAPII kinase activity [64]. Interestingly, one of the *S.pombe* PP1 catalytic subunits, Dis2, interacts with a PNUTS-like protein Ppn1. However, neither of the PP1 catalytic subunits (Dis2 and Sds21) modulate RNAPII-CTD Ser5 phosphorylation [65].

Furthermore, Ppn1 lacks all C-terminal RNA-binding motifs found in metazoan PNUTS proteins, which are required for its association with active RNAPII. These differences suggest that a regulatory module comprising PNUTS-PP1 and Cdk11 may have co-evolved after the divergence of yeast from a common ancestor. We note this coincides with the advent of more complex genomes containing a greater abundance of non-coding sequences, including intronic DNA, that by necessity requires greater transcriptional regulation.

Efforts are now directed towards identifying antagonistic kinases that oppose the effect of PNUTS-PP1 on substrates, such as Cdk11. The DFG+7 Serine in Cdk11 closely matches the consensus phosphorylation site for many CMGC-group serine/threonine protein kinases with roles in transcription or co-transcriptional mRNA processing, highlighting the scope for cross regulatory interactions. This includes Cdk12, a kinase associated with transcription elongation and pre-mRNA splicing [66, 67], which we found in our PNUTS co-precipitates. Understanding the functional interplay between pairs of validated kinase and phosphatase will help shed light on the broader regulatory phosphorylation networks governing gene expression.

### Potential roles of *Drosophila* Cdk11 in transcription and RNA processing

Cdk11 has known roles in transcription elongation and mRNA splicing. By analysing the distribution of intronic mRNA reads by RNA-Seq, we found that *cdk11* loss of function leads to increased intron retention, consistent with a conserved role for *Drosophila* Cdk11 in splicing. We are currently unable to conclude whether effects on splicing are direct or indirect. Co-transcriptional splicing accounts for the majority of all mRNA splicing in *Drosophila* [68] and humans [69, 70], and is associated with phosphorylated RNAPII-CTD-Ser5 [71, 72], which physically interacts with the catalytic spliceosome [73-75]. Consequently, the splicing defects we observed might be caused by altered RNAPII-spliceosome dynamics. In mammalian cells, paused RNAPII marked with RNAPII-CTD Ser5 has been observed to accumulate on exons that are subject to alternative splicing, particularly during exon inclusion [76]. This accumulation may mirror the 5’ capping checkpoint at nascent transcripts closer to the transcription start site, which also involves RNAPII-CTD Ser5 [77]. Notably, the disruption of RNAPII-CTD-Ser5 phosphorylation leads to intron retention, emphasising its critical role in alternative splicing [73]. However, human Cdk11 is also recognised to directly phosphorylate splicing factors and is involved in spliceosome activation [23-25, 27, 31]. Alternatively, therefore, the splicing defects we have observed in *cdk11^P^* mutant may be caused more directly by aberrant phosphorylation and activation of the spliceosome. Although we did not observe any difference in Sf3b1 phosphorylation at the residues we examined, alternative sites in the N-terminus of Sf3b1 and other splicing factors have been identified as Cdk11 substrates in other systems. A systematic analysis of *Drosophila* Cdk11 targets will be required to address this issue, along with measurements of their phosphorylation status and effects on splicing at specific gene loci after acutely disrupting cdk11 function.

### Regulatory roles of Cdk11 dephosphorylation

Building on our understanding of *Drosophila* Cdk11’s potential roles in transcription and RNA processing, we turned our attention to the regulatory roles of Cdk11 dephosphorylation using a *cdk11^SE^* phosphomimetic mutant. In contrast to the effect of *cdk11* loss of function, we observed the widespread loss of intronic snoRNAs in *cdk11^SE^* animals. The expression of snoRNAs and their host genes has been reported to involve multistep regulation, including transcription from dual-initiation promoters [78], Nonsense-Mediated Decay [79] and alternative snoRNA processing [80], with snoRNA stability and terminal stem formation also contributing to snoRNA expression expression levels [81]. In humans, intronic snoRNAs typically mature through splicing, debranching and exonucleolytic processing, or through an alternative pathway involving endonucleolytic cleavage [82]. *Drosophila* snoRNA processing is poorly described but is possible that dephosphorylation of Cdk11 is required for regulation of one or more analogous steps in this maturation pathway. Notably, we found that on a sample of *Drosophila* genes, RNAPII speed was reduced. It remains to be determined whether altered activity of RNAPII-CTD kinases or RNAPII speed in *cdk11^SE^* larvae is sufficient to account for the observed defects on snoRNAs. However, RNAPII speed has the potential to influence snoRNA splicing by modifying the interval between the formation of competing splice sites, influencing the recruitment of RNA-binding proteins, and altering the window of opportunity for transcript folding and covalent RNA modification. The underlying mechanisms will be worthy of attention given the emerging roles in snoRNAs in functions such as survival and longevity [83], which we found also to be disrupted in *cdk11^SE^*. Given the opposing effects of *cdk11^P^* and *cdk11^SE^* on the abundance of extragenic transcripts, another aspect that merits further examination, is the role of Cdk11 in PNUTS-mediated suppression of extragenic transcription [19, 20].

In summary, we identify here a stable regulatory complex that is comprised of PNUTS-PP1 and Cdk11. Within this complex, PNUTS-PP1 dephosphorylates a highly conserved Serine residue situated at the DFG+7 position within the activation loop of Cdk11, altering Cdk11 activity. We propose this may serve to regulate the dynamic interplay between transcription and RNA processing by modulating RNAPII-CTD-Ser5 phosphorylation. More broadly, it is noteworthy that phosphorylatable residues are present at the DFG+7 position in multiple cyclin dependent kinases (including Cdk7, 8, 11-13 and 19). Reversible phosphorylation at this position in the activation loop may therefore have emerged as a widely utilised biological strategy to modulate Cdk function.

## Methods

### Engineering of Drosophila strains

#### Overexpression constructs

UAS-PNUTS overexpression constructs were generated by cloning codon optimised full-length PNUTS coding sequences (GeneArt, Thermofisher) carrying synonymous RNAi-resistant mutations [84] into pPGW-attB (Megraw lab, Florida State University) using Gateway cloning to create an N-terminal GFP fusion. An HA tag (YPYDVPDYA) was incorporated at the C-terminus. *Genomic transgenes*: A 6kb genomic *cdk11* construct was made by cloning a synthetic DNA fragment encompassing the *cdk11* transcription unit (GeneArt, Thermofisher), -705bp of the ATG to +359bp of the stop codon, into the *Bam*HI/*Not*I sites of the fly transformation vector pW8-attB. The phosphorylation site in *cdk11* was mutagenised via Gibson cloning to generate pW8-attB::cdk11^S712A^ and pW8-attB::cdk11^S712E^. pW8-attB::cdk11 was digested with *Xba*I/*Bsi*WI and Gibson assembly using two fragments and overlapping primers containing the mutation of interest on pW8-attB::cdk11 template DNA.

#### InDisruPT

The construction of conditional alleles is to be described elsewhere. In brief, these consist of a modified version of a PNUTS genomic rescue construct [9], including attB site, FRT sites flanking the last coding exon, 3’ GFP tag and duplicated 3’ exon with an mCherry fusion. This was inserted on the third chromosome, combined with a *PNUTS* null allele, *PNUTS^13B^*, that lacks all coding sequence for *PNUTS*, and heat shock inducible FLP (*hsFLP*).

#### Transgenesis

PNUTS and cdk11 transgenic lines were made by PhiC31-mediated site-specific integration at the attP2 and attP40 landing sites, respectively. Transgenesis was performed by either the Cambridge or Manchester Fly Facilities (University of Cambridge, UK; University of Manchester, UK) or BestGene Inc (Chino Hills, USA).

#### Generation of cdk11 mutant alleles

Mutagenesis was performed using CRISPR/Cas9-mediated genome editing by homology-dependent repair (HDR) using a guide RNA and a dsDNA donor plasmid by Wellgenetics. In brief, the two homology arms of *pitslre*/*cdk11* were amplified by Phusion High-Fidelity DNA Polymerase (Thermo Scientific) from genomic DNA of the injection strain, and cloned together with a PBacDsRed cassette, containing 3X Pax3 and hsp70 promoter, DsRed2 and SV40 3’UTR (3xP3 DsRed), flanked by PiggyBac terminal repeats, into pUC57-Kan. gRNA sequences are available on request. Plasmids containing *pitslre*/*cdk11* gRNAs and hs-Cas9 were microinjected together with pUC57-Kan donor plasmid into *w^1118^* embryos. F1 flies with 3xP3 DsRed expression were examined by genomic PCR and sequencing to confirm insertion of the marked PBac into the *cdk11* coding sequence (*cdk11^P^*). After confirming transgene insertion, 3xP3-DsRed was excised by Cre-lox mediated excision and site-directed lesions confirmed by sequencing. A silent TTAA motif was introduced to the coding sequence, together with phosphomimetic mutation (S538/712E) carried on the homology arms, to generate the *cdk11^SE^* allele.

### Genotypes of strains

*<u>Figure 1, 2</u>*

*w^1118^*

*w^1118^; da-GAL4 , UAS-GFP-PNUTSWT-HA*

*w^1118^; da-GAL4 , UAS-GFP-PNUTS^W726A^-HA*

*Figure 3*

*hsFLP; PNUTS^13B^; PNUTSWTGFP::PNUTSWTmCherry/+*

*hsFLP; PNUTS^13B^; PNUTSWTGFP::PNUTS^W762A^mCherry/+ PNUTS^13B^;*

*PNUTSWTGFP::PNUTSWTmCherry/ovoFLP PNUTS^13B^;*

*PNUTSWTGFP::PNUTS^W762A^mCherry/ovoFLP*

*Figure 5*

*w^1118^*

*w^1118^;; cdk11^P^/ cdk11^P^*

*w^1118^; [cdk11^wt^]; cdk11^P^/ cdk11^P^*

*w^1118^; [cdk11^SA^]; cdk11^P^/ cdk11^P^*

*w^1118^; [cdk11^SE^]; cdk11^P^/ cdk11^P^*

*w^1118^;; cdk11^SE^ /cdk11^SE^*

*Figure 6,7*

*w^1118^*

*w^1118^;; cdk11^P^/ cdk11^P^*

*w^1118^;; cdk11^SE^ /cdk11^SE^*

### *Drosophila* culture and embryo collection

Fly stocks were kept at 18°C, 22°C or 25°C on standard yeast/dextrose ASG medium (1% (w/v) agar, 8.5% (w/v) dextrose, 2% (w/v) yeast, 6% (w/v) maize, 0.25% (v/v) nipagin, 0.3% (v/v) propionic acid). For embryo collections on apple juice agar, *en masse* crosses were performed in population cages at 25°C under a 12:12 hr light:dark cycle. For this, UAS-PNUTS lines were crossed to *da-GAL4* virgin females selected using a Y chromosome insertion of heat shock inducible cell lethal transgene, *hs-hid*. 0-18h embryos were dechorionated with bleach, washed with water and stored in Ultra High Recovery tubes (StarLab) at -80°C after being snap frozen in liquid nitrogen.

### Co-immunoprecipitation (Co-IP)

#### Embryo lysis

Embryos were homogenised (∼10 strokes) in cooled 1 ml Glass dounce homogenisers (Glass precision engineering) in 500 µl chilled lysis buffer (100 mM Tris-HCl pH 7.6, 150 mM NaCl, 0.5 mM EDTA, 0.5 % Triton-X, 0.5 % NP-40 and protease inhibitors) with Benzonase nuclease 1:250 (25 U/µl, Merck Millipore). Lysates were incubated on ice for 30 min, homogenising with 1-2 strokes every 10 min before being centrifuged at 7800 xg for 10 min at 4°C. Total protein concentration was analysed via nanodrop and normalised across samples to standardise IP input. *Anti-HA co-immunoprecipitation (Co-IP)*: Unless otherwise stated, all steps were performed on ice in the presence of protease/phosphatase inhibitors (Mini EDTA-free protease inhibitor cocktail (Roche) and 1x PhosSTOP (Roche) to minimise protein degradation; 1.5 ml Ultra High Recovery eppendorf tubes (Star Lab) were used to minimise sample loss. 15 µl per IP of washed anti-HA magnetic beads (Pierce, ThermoFisher Scientific) were incubated with clarified lysate at 4°C overnight with end-over-end mixing. Beads were washed 3-6 times, for 0-2 minutes per wash, with HA IP wash buffer (40 mM Tris-HCl pH 8, 0.5 mM NaCl, 0.1 % NP-40, 1 mM EGTA, 6 mM DTT, and protease inhibitors) then once with HPLC water.

### Sample preparation for Mass spectrometry with dimethyl labelling

Following Co-IP, anti-HA beads were washed 3 times with 100 mM Triethylammonium bicarbonate (TEAB) then resuspended in 0.06% (w/v) Rapigest SF Surfactant (Waters) in TEAB for in-solution tryptic digestion and boiled at 80°C for 10 minutes to elute proteins. Dimethyl labelling was performed by in-solution labelling [35] except volumes of all reagents were increased 4-fold to account for the increased protein concentration. IPs were differentially dimethyl labelled, intermediate (with CH_2_O) for PNUTS^WT^ and light (with CD_2_O) for PNUTS^W726A^. Post-labelling, samples were combined 1:1 and desalted using Macro C18 spin columns (Harvard Apparatus). Peptides were eluted in ACN:H_2_O (80:20) with 1% Trifluoroacetic acid, centrifuged for 10 minutes at 1100 xg and dried to completion by vacuum centrifugation. Samples were subjected to strong cation exchange using 47 mm Cation Exchange discs (Empore Supelco) to remove Polyethylene glycol contamination originating from stored embryos. 10% of the digested material was used to analyse the PNUTS-interactome without further enrichment. The remaining 90% was enriched for phosphopeptides using Titanium dioxide (TiO_2_). For this, peptides were reconstituted in Loading buffer (80% ACN, 5% TFA with 1M glycolic acid) at 1 mg/ml and sonicated for 10 min in a water bath prior to incubation with TiO_2_ resin (GL Sciences; 1 mg/200 µg total protein). After washing with Loading buffer, 80% ACN, 1% TFA then 10% ACN, 0.2% TFA, peptides were eluted from the resin with ammonium hydroxide. Elutions were combined and dried to completion in a vacuum centrifuge. For LC-MS/MS analysis, samples were reconstituted in H_2_O:ACN (97:3) with 0.1% formic acid and sonicated 10 min in a water bath.

### Mass spectrometry data acquisition and data analysis

Peptides were separated by reversed-phase capillary HPLC on an UltiMate 3000 nano system (Dionex) coupled in-line to an Orbitrap Fusion mass spectrometer (Thermo Scientific, Bremen, Germany). Peptides were loaded onto the trapping column (PepMap100, C18, 300 µm x 5 mm) with 2% (v/v) ACN, 0.1% (v/v) TFA using partial loop injection at a flow rate of 9 µl/min for seven min, then resolved on an analytical column (Easy-Spray C18 75 µm x 500 mm, 2 µm bead diameter column) at a flow rate of 0.3 µl/min using either a 60 min gradient (for analysis of TiO2 enriched samples) or 120 minute gradient (for analysis of unenriched samples) of A (0.1% Formic acid) to B (80% ACN with 0.1% Formic acid), starting at 96.2% A and ending at 50% B. MS(/MS) data were acquired as [85]. In brief, for unenriched IP samples; MS1 spectra were acquired over m/z 350-2000 in the Orbitrap (120K resolution at m/z 200) with AGC set to accumulate 2E5 ions, with a maximum injection time of 50 ms. MS2 data was acquired in a data-dependent manner using a top speed approach (cycle time of 3 s) with HCD fragmentation in the ion trap (normalized collision energy set to 32%). The ion trap was operated in rapid mode with a maximum injection time of 35 ms, target value of 1E4 and a fixed first m/z of 110. The intensity threshold for fragmentation was set to 5K and included charge states from 2+ to 5+. A dynamic exclusion window of 60 s was applied with a mass tolerance of 0.5 Da.

For TiO2 enriched IP samples, MS1 spectra were acquired over m/z 350-2000 in the Orbitrap (60K resolution at m/z 200) with AGC set to accumulate 2E5 ions, with a maximum injection time of 50 ms. MS2 data was acquired in a data-dependent manner using a top speed (cycle time of 3 s) approach with HCD fragmentation (normalized collision energy set to 32%) in the Orbitrap at 30K resolution with a maximum injection time of 100 ms, target value of 5E4 and a fixed first m/z of 110. The intensity threshold for fragmentation was set to 50K and included charge states from 2+ to 5+. A dynamic exclusion window of 60 s was applied with a mass tolerance of 10 ppm.

For quantification of dimethyl labelled samples, data were processed using MaxQuant (version 1.6.0.16, Max Planck Institute of Biochemistry) and used to search Uniprot and UniDrosophila database with the Andromeda search engine allowing up to 2 missed cleavages and 1% false discovery rate. Perseus (v. 1.6.1.1) was used to remove hits to the contamination database. Since PNUTS was GFP-tagged, we also removed proteins that precipitated with GFP alone with GFP-Trap. Selecting high-confidence interactors based on prevalence: we selected proteins identified in 2 or more PNUTS IPs, and then excluded proteins identified in 2 or more *w^1118^* control IPs. Abundance-based interactor selection: For each protein identified in 2 or more PNUTS IPs, we used EmPAI (Exponentially modified protein abundance index) values derived from the number of identified peptides divided by the theoretical number of observable peptides expected for any given protein, as a measure of relative abundance (Ishihama et al., 2005) in w^1118^ control IP samples relative to PNUTS IP. Any proteins <2-fold enriched in PNUTS IPs compared to w^1118^ control IPs were considered to be non-specific binders. Filtered data was imported into R for plotting and statistical analysis using the limma package for Bayesian analysis [86]. String DB/Cytoscape String (v. 10.5, accessed at https://string-db.org) was used for network analysis. Only high (>0.7) or medium (>0.4) confidence interactions were included, with no additional white nodes. MCL clustering was used to identify clusters of shared function to colour nodes. Cytoscape (v. 3.5.1) was used to rearrange and customise String network nodes. Gene ontology analysis and functional annotation was performed using the Database for Annotation, Visualization and Integrated Discovery (DAVID).

### Antibody production and validation

Custom rabbit polyclonal antibodies against *Drosophila* Cdk11 phosphorylated at S538/712 were raised by Pierce Custom Antibodies using the following peptide coupled to keyhole limpet haemocyanin: C-LAREYG(pS)PIKKYT-amide. pCDK11 antibody was subjected to two rounds of affinity purification positive selection using the phosphorylated peptide and negative depletion against a non-phosphorylated control peptide. Specificity of the phospho-antibody was validated using lysates of second instar larval extracts overexpressing UAS-cdk11 (UAS-pitslre, RRID:BDSC_55066) or dsRNA from an inverted repeat construct UAS-cdk11^IR^ (TRiP.HMS02389), under the control of *da-Gal4.* Antibodies against *Drosophila* Sf3b1 phosphorylated at Ser137 were generated in the same way using the following peptide for immunisation: CRRHIII(pS)PERADP-amide. Sf3b1 antibodies were tested using UAS-sf3b1^IR^ (TRiP.HM05176).

### Imaging acquisition and analysis

Adult females of the desired genotypes were fattened on yeast paste for 2 days at 25°C prior to dissection in 1x PBS. Ovaries were fixed and stained as [87]. In brief, tissues were fixed in 3.7% PFA for 20 minutes, washed with PBS plus Tween 20 (PBST; 1× PBS, 0.2% Tween 20) and blocked with PBST and FBS (PBTB; 1× PBS, 0.2% Tween 20, 5% FBS) for 1-2 hours at room temperature. Ovaries were incubated with anti-pCDK11 antibody (1:50; this study) in PBTB overnight at 4°C. Then, ovaries were washed with PBST three times 15 min and incubated with a AlexaFluor(AF)405-conjugated secondary antibody (1:500; Life Technologies) overnight at 4°C in PBTB. After final washes with PBST, samples were incubated with TO-PRO-3 Iodide (1:1000, Life Technologies) in 1x PBS for 20 minutes and then mounted with VectaShield mounting media (Vector Laboratories). Slides were imaged using an LSM 780, or LSM880 confocal microscope (Carl Zeiss) using a 40x oil objective with excitation of AF405, GFP, mCherry and TO-PRO-3 Iodide using 405nm, 488 nm, 561 nm and 633 nm laser lines, respectively. Quantification of fluorescent signals was performed using linescans in FIJI/ImageJ (https://fiji.sc/). The profiles of signal intensities were displayed in two-dimensional graphs generated in Graphpad Prism.

### Recombinant proteins and enzyme assays

The full-length coding sequences for *Drosophila* Cdk11 (as overexpression constructs above) and Cyclin L/CG16903 were synthetised by GeneArt and cloned into a proprietary N-terminal His tag vector for co-expression in Sf9 insect cells by SignalChem Biotech (Richmond, Canada). After 72 h expression, cell lyasates were harvested and incubated with Nickel beads at 4°C, washed with 10 mM imidazole containing wash buffer (50mM Na2HPO4 PH7.5, and 300mM NaCl), and eluted with a gradient of 10-200 mM imidazole in wash buffer supplemented with 0.25 mM DTT. Purity of recombinant Cdk11/Cyclin L was determined to be ∼70% by densitometry of Coomassie-stained gels, with the recombinant proteins having an apparent molecular mass of 111 kDa and 63 kDa, respectively. Kinase assays were performed *in vitro* by incubating 100 ng/µl recombinant human RNAPII-CTD (Abcam, ab81834, >95% purity) for 15 min at 30°C in the presence of 50 ng/µl CDK11s/Cyclin L, 100 µM ATP, 50 mM Tris-HCl (pH 7.4), 100mM NaCl, 1 mM DTT, 100 mM MgCl_2_, and immunoblotted using total RNAPII-CTD (Chromotek, #4H8, 1:1000) and phospho-specific RNAPII-CTD antibodies: phosho-Ser 2, 5, 7, Thr4 (Chapman et al., 2007; Chromotek). For kinase assays with Sf3b1, residues 1-463 were amplified from a human RPE1 cDNA library and cloned into pGEX-GP-1, prior to expression as a GST-fusion in T7 Express cells (New England Biolabs). Recombinant human Sf3b1 was purified with a GS Trap (Cytiva). GST-Sf3b1 was used as a substrate in Cdk11/Cyclin L assays as above, except that incubations were for 60 min and were followed by immunoblotting with phospho-Sf3b1 (Thr 313) (Cell Signalling, #D8D8V, 1:1000) and total Sf3b1 (MBL, anti-Sap 155 mAb, #D221-3, 1:1000) antibodies. For kinetic experiments, we synthesised a CTD peptide conjugated to an N-terminal FAM fluorophore (5-FAM-(YSPTSPS)_3_-CONH_2_, Pepceuticals) and monitored CTD peptide (2 µM) phosphorylation in real-time in the presence or absence of 1 µg Cdk11 (WT, S712E, or S712A)/Cyclin L and 0.5 µg CDK7/cyclin H (SignalChem Biotech Inc) using a LabChip EZ Reader microfluidic platform (PerkinElmer) [88]. Assays were performed in the presence of 1 mM ATP, 50 mM Hepes (pH 7.4), 0.015% (v/v) Brij-35, 1 mM DTT, and 5 mM MgCl_2_. Pressure and voltage settings were −1.0 psi and −1800 V (upstream voltage) and −500 V (downstream voltage), respectively. Peptide phosphorylation kinetics (with substrate phosphorylation limited to <10%) were shown as the ratio of phosphopeptide:peptide present in each sample of a 384-well plate. For Western dot blots, Cdk11/Cyclin L and CDK7/cyclin H and 5-FAM-(YSPTSPS)_3_-CONH_2_ were incubated overnight at 25°C in the presence of 1 mM ATP and 5 mM MgCl_2_, transferred onto nitrocellulose membranes and blotted with RNAPII-CTD antibodies (as above). For immunoblotting assays of Cdk11-S712 phosphorylation, we employed anti-pCdk11 (custom antibody, see above, 1:1000), anti-GFP (Life Technologies, #A111122, 1:2000), anti-RFP (Chromotek, #6G6, 1:1000) and anti-total Cdk11 ([89], a gift from Nic Tapon) antibodies.

### RNA-sequencing and data analysis

RNA samples were extracted from homozygous c*dk11^PBac^*and Cdk11^S712E^ first and second instar larvae, along with matching developmentally-staged *w^1118^* controls, using TRIzol (Thermofisher) in triplicate. The quality of RNA samples was tested using Tape station 4.1.1 (Agilent Technologies) and ribosomal RNAs (rRNA) were depleted using RiboZero kit (Illumina). RNA libraries were prepared using NEB E7760 Directional library prep. RNAs were sequenced to a read depth of 15–30 million reads on the Illumina platform using NovaSeq (Novogene). Raw data (fastq) files were processed with trimgalore, a wrapper script, to automate quality and adapter trimming. Reads were filtered to remove rRNA with Bowtie 2 mapping [90]. Then the data was analysed using two pathways. First, we used Salmon, which corrects for fragment GC-content bias, to numerate the expression of transcripts [91]; differential gene expression was quantified using DESeq2 [92]. Gene Ontology enrichment was performed using Metascape (https://metascape.org/gp/index.html)[93]. Secondly, we used HISAT2 (hierarchical indexing for spliced alignment of transcripts 2) [94] for graph-based alignment to the *Drosophila melanogaster* genome (BDGP6.32) and analysed changes in alternative splicing (AS), particularly intron retention, using NxtIRF [57, 58].

Complete-linkage hierarchical clustering analysis based on Euclidean distances was performed to assess the relationship between all samples and the reproducibility of biological replicates. Normalised data from regularized log transformation (rlog) of raw gene-level count data by DESeq2 was used as input for the clustering. We identified the percentage of reads aligning to intergenic sequences when genome-mapping with HISAT2 using the CollectRnaSeqMetrics program from the Picard suite of tools (Broad Institute, 2022). Aberrant gene transcription events, including those that fail to terminate correctly (readthrough), those that are aberrantly transcribed due to failure of upstream termination (read-in), and novel transcription of downstream regions arising from readthrough (downstream of gene transcription (DoG), were characterized by analysis of HISAT2 mapped data using ARTDeco with its default settings [61]. ARTDeco quantifies read-in and readthrough transcription as summary statistics based on the ratio of the expression levels of these regions compared to the expression level of the gene region itself for the top 1000 most highly expressed genes. The medians of the read-in and readthrough level distributions for the top-expressed genes represent useful summary statistics for characterizing the overall degree of read-in or readthrough in a given sample [61]. DoG transcription is identified by quantifying reads mapping to regions downstream of genes using a 500 bp rolling window approach relative to the expression of the upstream gene. The median FPKM expression of all DoGs identified is used as a summary statistic for the global level of DoG transcription in each sample.

To estimate transcription speeds, we followed the method of Debes et al 2023 [95], using scripts modified from those available at https://github.com/beyergroup/ElongationRate, run in R version 4.3.2. Briefly, this method uses coverage of introns from RNA-Seq reads obtained from rRNA depleted libraries to estimate transcription speed from the slope of a linear model fit to coverage depth over position. In brief, we aligned RNA-Seq reads to the *Drosophila melanogaster* reference genome version BDGP6.46 using STAR version 2.7.11a [96], as Debes et al. 2023 except with reduced stringent filtering of reads. Introns annotated by STAR that passed thresholds of at least 4 uniquely mapped reads crossing the intron junction and at least one alignment overhanging the splice junction by at least 10 bases, resulted in 13204 and 17784 introns retained for cdk11^SE^ and cdk11^P^ comparisons, respectively. We then estimated the coverage slope for introns mapping to major chromosome arms, using coverage of uniquely mapped reads estimated by STAR to estimate the slope using the lm function in R. For each mutant and control comparison, the slopes files for samples were merged and filtered for introns where the slopes were negative for each sample, resulting in 415 and 403 intron comparisons for cdk11^SE^ and cdk11^P^ comparisons, respectively.

### Database submissions

RNA sequencing data are available from NCBI Gene Expression Omnibus (https://www.ncbi.nlm.nih.gov/geo) under accession number GSE256082.

Mass spectrometry proteomics data have been deposited to the ProteomeXchange Consortium via the PRIDE [97] partner repository (https://www.ebi.ac.uk/pride/) with the dataset identifier PXD051585.

## Acknowledgements

We wish to thank Timothy Megraw (Florida State University) and Nic Tapon (The Crick) for essential reagents. We would also like to thank the Centre for Proteome Research and the Centre for Cell Imaging, University of Liverpool, the Brighton Integrative Genomics Unit, University of Brighton, and the Manchester Fly Facility and the Gene Editing Unit, University of Manchester for research support. A.E.C was funded by the BBSRC (ref. 1510556), A.A.A. by the Saudi Arabia Cultural Embassy, and T.Z. by a Presidential Research Fellowship, University of Manchester. The work was supported by Medical Research Council (MR/K015931/1).

The authors declare no competing financial interests.

## Author contributions

Campbell AE: investigation, methodology, validation, formal analysis, visualisation, writing — review and editing. § These authors contributed equally.

Aljabri AA: investigation, methodology, validation, formal analysis, visualisation, writing — review and editing. § These authors contributed equally.

Hesketh A: investigation, formal analysis, visualization, data curation, writing — review and editing.

Byrne DP: investigation, supervision, formal analysis, writing — review and editing.

Bennett H: resources, writing — review and editing. Patel S: resources, writing — review and editing.

Brownridge P: supervision, formal analysis.

Zacharchenko T: resources, writing — review and editing.

Bucca G: resources, writing — review and editing.

Eyers PA: supervision, resources, writing — review and editing.

Betancourt AJ: supervision, methodology, formal analysis, writing — review and editing.

Eyers CE: supervision, resources, methodology, formal analysis, data curation, writing — review and editing.

Bennett D: conceptualisation, project administration, supervision, resources, methodology, formal analysis, visualisation, data curation, writing — original draft, writing — review and editing.

## Notes

### Competing Interest Statement

The authors have declared no competing interest.

